# Chromatin profiling data indicate regulatory mechanisms for differentiation during development in the acoel *Hofstenia miamia*

**DOI:** 10.1101/2023.12.05.570175

**Authors:** Paul Bump, Kaitlyn Loubet-Senear, Sarah Arnold, Mansi Srivastava

## Abstract

Chromatin profiling data can corroborate and generate hypotheses for regulatory events that underlie the control of gene expression in any biological process. Here, we applied the Assay for Transposase Accessible Chromatin (ATAC) sequencing to build a catalog of putative regulatory DNA during the process of embryonic development in an acoel. Acoels represent an enigmatic phylum-level lineage of animals, the Xenacoelomorpha, which is placed either as a sister-group to all other animals with bilateral symmetry or as an early-diverging ambulacrarian, two alternative phylogenetic placements that both position acoels equally well to inform the evolution of developmental mechanisms. We focused on the acoel *Hofstenia miamia*, a.k.a. the three-banded panther worm, which has emerged as a new laboratory research organism for whole-body regeneration that also enables the study of development from zygote to hatching. We profiled chromatin landscapes over a time course encompassing many major morphological events, including gastrulation, axial patterning, and differentiation of tissues such as epidermis and muscle. Broad patterns of chromatin accessibility and predicted binding of various transcription factor (TF) motifs identified major biological processes and their putative regulators, and we noted that differential accessibility tended to precede major developmental transitions in embryogenesis. Focused analysis of TF binding combined with single-cell RNA-seq data provided regulatory linkages for genes in a previously hypothesized differentiation trajectory for epidermis and generated new hypotheses for gene regulatory networks associated with the formation of muscle. This work provides a platform for the identification of developmental mechanisms in *Hofstenia* and enables comparisons of embryogenesis in acoels to other animals as well as comparisons of embryogenesis to regeneration.

## INTRODUCTION

Dynamic regulation of chromatin such that different regions of the genome become accessible over time is a measurable pattern that accompanies the regulation of gene expression during embryonic development and is crucial for establishing the animal body plan^1^. Chromatin dynamics during development have been studied in several systems, revealing key features of developmental gene regulation^2–5^. In zebrafish, promoters and enhancers increase in accessibility during zygotic genome activation, and specific maternally-loaded transcription factors prime genes for activation^4^. Consideration of three sea urchin species across different developmental modes demonstrated that global chromatin dynamics contribute to embryonic gene expression and can help explain the evolution of variable developmental life history strategies^5^. In the amphipod crustacean *Parhyale hawaiensis,* a chromatin analysis through embryogenesis revealed that the majority of regulatory regions in the genome had dynamic accessibility across development^2^. In the amphioxus *Branchiostoma floridae* the developmental trajectory of different cell lineages and underlying regulatory modules have been clarified with a combination of gene expression and chromatin state data^3^. These data, all derived from bilaterian species, are highly effective for advancing insights into species-specific biology. To broaden the taxonomic sampling for such data, and to glean shared properties of embryonic gene regulation, we focused on the animal phylum Xenacoelmorpha, a group that is highly informative for understanding the evolution of development given its placement either as the sister-group to all other bilaterians or to ambulacrarians^6–8^ (Figure 1A).

**Figure 1.**
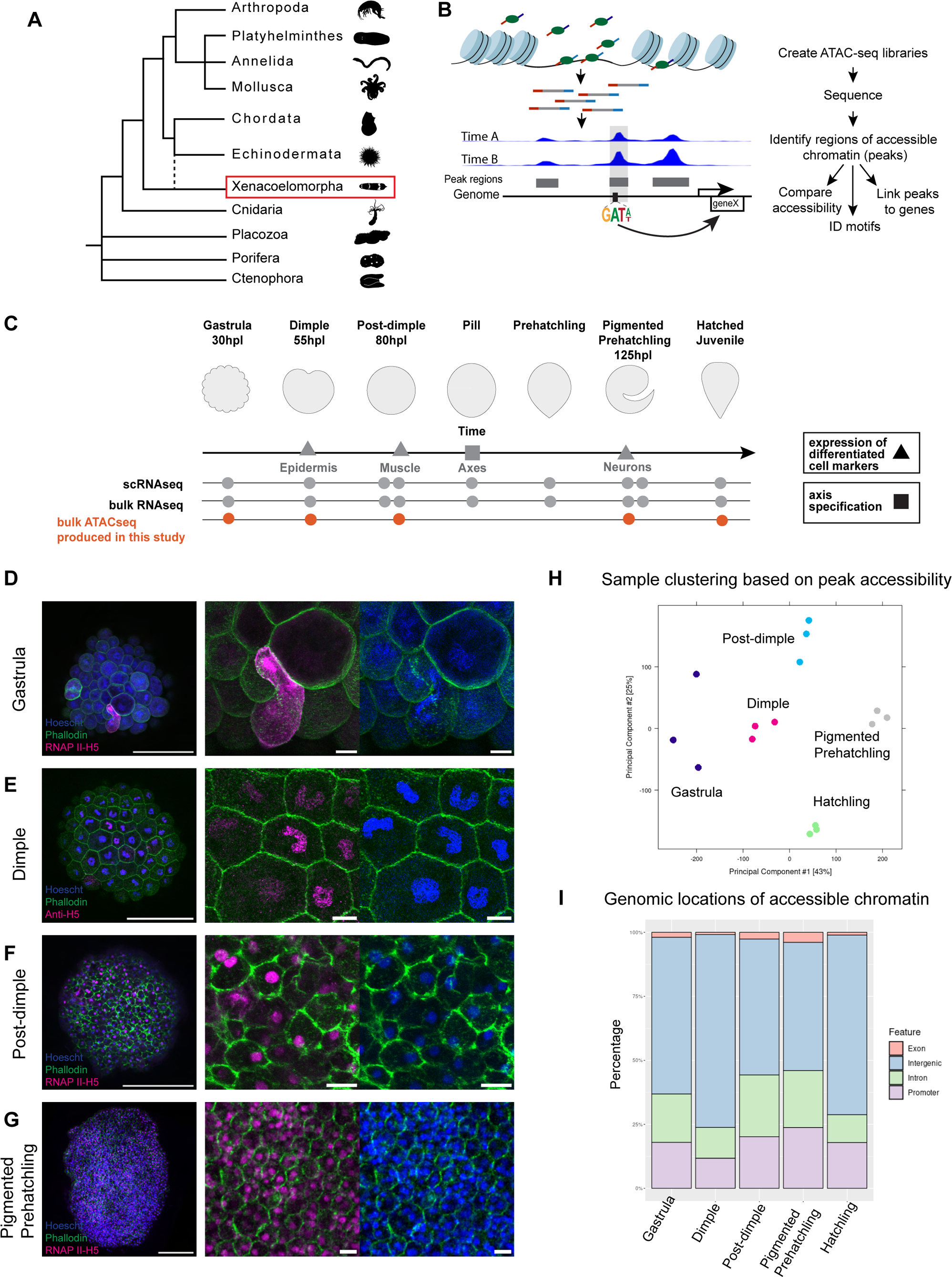
Epigenomic analysis of *Hofstenia miamia* development identifies regions of accessible chromatin. (A) Schematic phylogenetic tree of select metazoan lineages demonstrating the placement of Xenacoelomorpha (red box) as the likely sister group to all other bilaterians or sister to the Ambulacraria (dotted line). Public domain silhouettes courtesy of PhyloPic. (B) Schematic of chromatin profiling workflow illustrating how data was generated and regions of the genome are compared to each other. (C) Schematic of the different developmental stages that were sampled highlighting when differentiated cell markers are present as well as other previously published RNAseq that was analyzed in this study^9,14^. (D-G) Immunostaining of embryonic stages sampled: gastrula, dimple, post-dimple and pigmented prehatchling with RNAPII H5 antibody, BODIPY Phallacidin, and Hoescht. (H) Principle component analysis of triplicate ATAC-seq libraries that were generated in this study. (I) Bar plots showing the proportions of peaks contained within promoters, exons, introns, and intergenic regions across the different developmental timepoints.

The acoel *Hofstenia miamia*, also referred to as the three-banded panther worm, has emerged as a new xenacoelomorph model system. These animals are easily cultured in the laboratory, where they produce plentiful embryos that are accessible from the zygote stage to hatching^9^. *Hofstenia* hatched juvenile and adult worms show robust whole-body regeneration^10^ and previously, the Assay for Transposase Accessible Chromatin following by sequencing (ATAC-seq) was used in *Hofstenia* to identify regulatory regions associated with regenerative processes, as well as to connect these regions with transcription factor binding and to elucidate genetic factors important for reshaping the regenerative chromatin landscape^11^. Along with a recently updated genome, these analyses have been leveraged to understand *Hofstenia* wound-induced gene regulatory networks,^11^ patterning after wounding,^12^ and the specification of neurons in a wound response context^13^. We reasoned that, similarly, investigating regions of accessible chromatin and regulatory landscapes throughout *Hofstenia* embryogenesis could provide insight into how the epigenome changes and which regulatory decisions might underlie major developmental processes.

Here we utilized ATAC-seq to understand how different regions of the genome are organized during development (Figure 1B). ATAC-seq of developing embryos revealed that the epigenome is highly dynamic during development and that accessibility of cohorts of genomic regions correspond to major developmental transitions. Additionally, we find that the predicted binding of certain transcription factor motifs that are variably accessible during development can point to regulatory linkages for the known differentiation trajectories, such as for the epidermis, as well as provide novel insights into transcriptional decision-making for other tissues such as the muscle.

## RESULTS

### Epigenome analysis of *Hofstenia miamia* development identifies regions of accessible chromatin

To understand how the epigenome changes across *Hofstenia* development we sampled several key time points that were previously characterized and subjected to both bulk and single-cell RNA sequencing (Figure 1C)^9,14^. *Hofstenia* development begins with a series of duet cleavages, after which the larger macromeres of the embryo are internalized. This results in the gastrula stage, the first time point we sampled. At this developmental time point there is no strong transcriptional evidence of cell lineage markers^9^. To investigate whether this might indicate that this stage occurs prior to the start of zygotic genome activation, we used an RNAPII-H5 antibody which labels a phosphorylated epitope on the carboxy-terminal domain of the large subunit of RNA Polymerase II (RNAP II) and is linked with the process of transcriptional elongation^15^. We detected only one or two RNAPII-H5 positive cells per embryo, indicating the potential start of minor zygotic genome activation (Figure 1D). Following additional cell divisions, the dimple stage occurs when a ring of actin becomes restricted to one side of the embryo, and a second cell internalization event takes place. The dimple stage also corresponds to the first evidence of tissue differentiation based on single cell RNA-seq (scRNA-seq), where transcription factors (TFs) as well as some differentiated markers corresponding to the epidermis are expressed^14^. At this stage we detected RNAPII-H5 in multiple cells of the embryo, suggesting that the zygotic genome is now active (Figure 1E). This is consistent with our expectation that transcription should be active in many cells if they are undergoing differentiation. Next, in the post-dimple stage, the embryo appears smooth, and although no major morphological events corresponding to the adult body plan are yet observable, various molecular trajectories for tissue types such as muscle and gut can be identified and in addition, molecular markers of when the main body axes are detectable^9^. At this stage RNAPII-H5 was high across the embryo (Figure 1F). The last embryonic stage sampled was the pigmented prehatchling (Figure 1G) in which pigment granules have begun to form, the worm has begun to develop a classic vermiform shape with a pointed tail in the posterior, and transcriptional differences between the major tissue types in *Hofstenia* are present. Finally, we sampled a hatched juvenile worm which has emerged from the eggshell and has begun to swim and crawl.

After preparing pooled embryos from each developmental stage for ATAC-seq to recover between 50,000-100,000 cells per sample, cells were lysed, nuclei were transposed, and sequencing libraries were amplified. After sequencing, demultiplexing and barcode adaptor removal, reads were mapped to the *Hofstenia* genome, quality control was performed (see Methods, Supplement 1), and duplicate reads were removed. We utilized Genrich replicate peak calling, which first analyzes each sample individually and calculates *p-*values which are then combined with Fisher’s method into a new corresponding *q-*value^16^. Each sample was loaded into DiffBind with its corresponding peak file to generate a new consensus file of any peak, which represents regions of accessible chromatin, that was called in at least two samples. This consensus peak file allowed us to identify the regions of the genome that these peaks correspond to called “peak regions”. These can be further analyzed in downstream pipelines for additional genomic features associated with accessible regions of chromatin, such as predicted transcription factor binding sites. Principle component analysis revealed that replicates from a given time point clustered together (Figure 1H), a result also reflected in hierarchical clustering (Supplement 3A).

To understand how the accessibility of different features of the genome changed across development we determined which peaks occurred in either the promoter, intron, exon, or intergenic regions of the genome and asked how these proportions of accessible regions change across development (Figure 1I). In accordance with ATAC-seq studies of regenerating tissue in *Hofstenia*, we found that a substantial portion of the accessible genome is found within intergenic regions^11^. We noted that the proportion of peaks, or “open” chromatin, in the promoter was diminished at the dimple and hatched juvenile stages, relative to the proportion detectable in the pigmented prehatchling (Figure 1I). These patterns based on the proportion of accessible genome also corresponded with the pattern observed in the total number of peaks called at each timepoint (Supplement 3B), which also showed a reduction in the number of peaks at dimple and hatched-juvenile stages.

### Differential accessibility across the *Hofstenia* genome reflects major developmental transitions

Using the program DiffBind to compare chromatin accessibility throughout *Hofstenia* development, we identified regions of the genome that opened or closed throughout development or between adjacent timepoints (see Methods). We identified ∼39k peaks total across all timepoints, reflecting a catalog of putative regulatory regions that could operate during the developmental stages we sampled. Of these, ∼35k regions showed statistically significant dynamic changes in chromatin accessibility (adjusted p <0.05,Wald test). Next, we performed hierarchical clustering on this dynamic peakset, which enabled us to identify major patterns in these dynamics (Figure 2A, Supplement 3D). Notably, while there was a large number of accessible peaks present at the gastrula stage, very few of these peaks exhibited increased accessibility at the dimple stage and in fact, many displayed decreased accessibility. As development progressed through post-dimple and pigmented prehatching, the number of newly emerging dynamic peaks increased until hatching, when chromatin regions tended to close.

**Figure 2.**
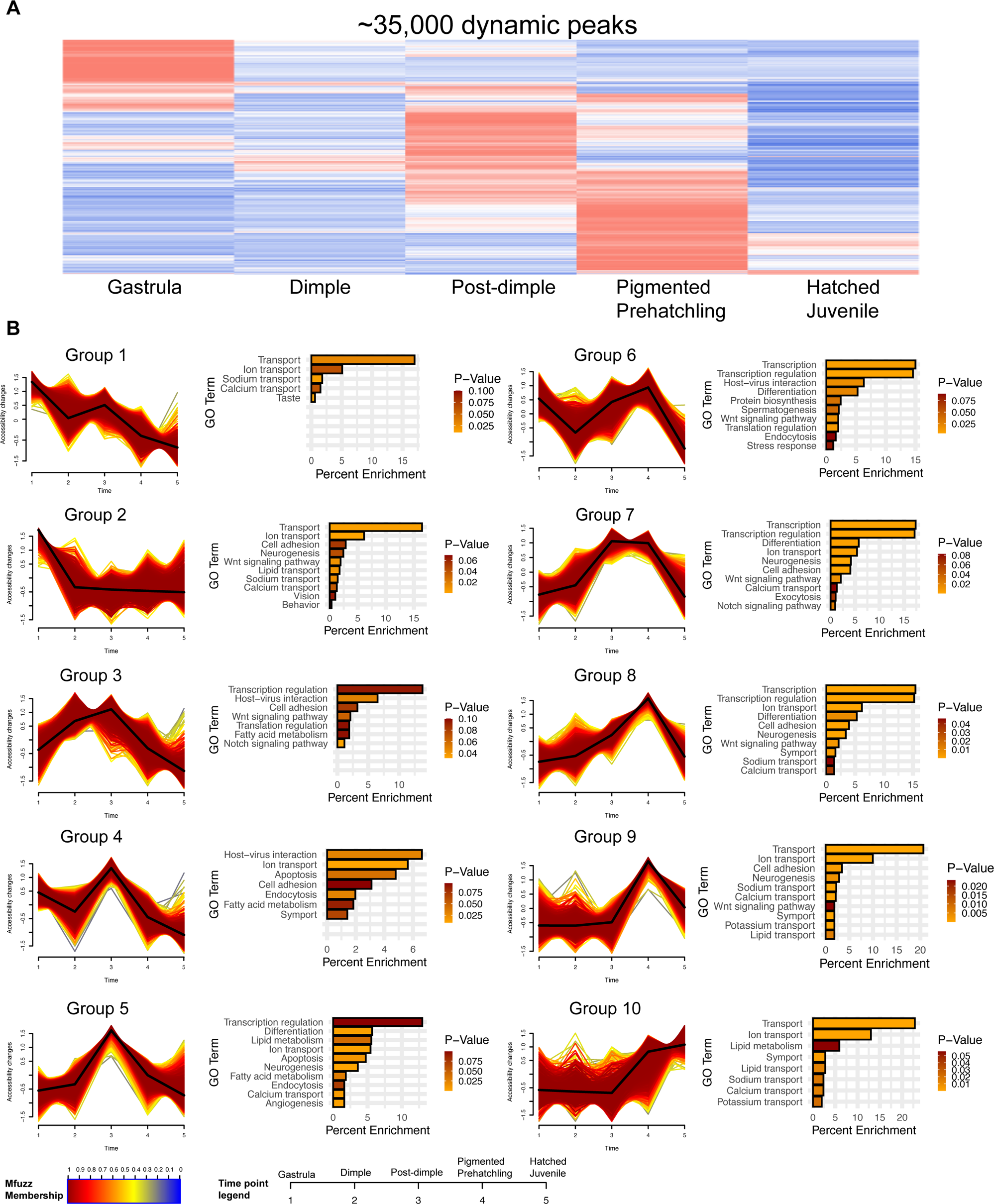
Differential accessibility of the Hofstenia Genome Reflects Major Developmental Transitions. (A) Heatmap of all chromatin peaks that show significant changes during Hofstenia development. (B) Fuzzy c-means clustering of regions of accessible chromatin with each cluster plotted individually. GO Functional Annotation enrichment for different peaksets, x-axis denotes the strength of GO Term enrichment and bars are colored by p-value.

To quantitatively group dynamic peaks based on their overall behavior in the developmental time course we sampled, we performed fuzzy c-means clustering^17^ and recovered several major trends (Figure 2B). These included sets of regions with highest accessibility at gastrula (Groups 1 and 2), regions with highest accessibility at hatchling (Group 10), regions with low accessibility at gastrula and hatched juvenile, but with high accessibility at an intermediate time point (Groups 3, 5, 7, 8, 9), and regions with low accessibility at dimple and hatched juvenile but with high accessibility at an intermediate time point (Groups 4, 6).

To understand which biological processes might be associated with these regions of increased chromatin accessibility, we first linked these peaks to genes using the peak annotation tool UROPA, and then used Gene Ontology (GO) analysis to ask whether these cohorts of genes were enriched for specific biological functions^18,19^. For peaks that showed greatest accessibility during gastrula (Groups 1 and 2) we found enrichment for GO terms relating to cellular transport-associated categories including ion, sodium, and calcium transport, indicating that membrane transport processes could play important roles early in *Hofstenia* development. In the set of peak regions with highest accessibility at the dimple (Group 3) we found the first instance of transcription-related terms (i.e., transcription regulation and transcription), concordant with our observation of RNA polymerase II activity in Figure 1D. Many other transcription related terms are also common in other groups that exhibited highest peak accessibility at later time points, such as the post-dimple and pigmented prehatchling (Groups 5, 6, 7, 8). When we asked specifically which genes populated these GO categories, we noted many of them corresponded to transcription factors (TFs), consistent with differentiation of tissue lineages at these timepoints^9^. Some of the terms in these categories were shared between clusters, potentially pointing to multiple roles of a single gene during development, but others were unique to one cluster. To gain further resolution at what the roles of some of these TFs might have at different times, we decided to compare two clusters comprised of peaks that both had high accessibility late (Clusters 6 and 7), but only one of which also had high accessibility early, at gastrula (Cluster 6). We found ninety-two genes shared by both Cluster 6 and Cluster 7 attributed to the transcription term, fifty unique genes in Cluster 6 which included terms related to chromatin remodeling, and fifty-six unique genes in Cluster 7 which included terms related to differentiation and notch signaling (Supplement 3C). This suggests that at late timepoints, differentiation and notch signaling may be important, consistent with neurogenesis or other tissue differentiation processes occurring. Chromatin remodeling, however, likely plays a role not just at this late stage, but also may be involved in early developmental processes. Cell adhesion appeared in clusters with peaks that had high accessibility at a number of timepoints (Groups 2-4, 7-9), pointing to the important role of cell-cell interactions in properly creating an animal body plan. Finally, Wnt signaling was enriched in multiple groups that had the most accessible peaks at a variety of time points (Groups 2,3, 6-9), suggesting Wnts may be important in multiple facets of development.

To complement the fuzzy c-means clustering analyses, which groups peaks based on patterns of opening and closing over time, we also sought to identify peaks that changed significantly between consecutive stages to understand the sequential developmental events that occur across *Hofstenia* embryogenesis. To do this, we used DiffBind to perform pairwise comparisons to identify the most differentially accessible peaks between adjacent timepoints. Consistent with the overall decrease in peak accessibility at the dimple stage we had previously observed (Figure 2A), we saw fewer peaks with greater accessibility at the dimple compared to the gastrula and post-dimple stages relative to comparisons between the post-dimple and pigmented prehatchling (Supplement 4A).

In general, the patterns identified with hierarchical clustering, fuzzy c-means clustering and differential accessibility all matched previous bulk RNAseq data in *Hofstenia* development which suggested some of the early molecular events are related to cell movement, followed by transcription, and eventually categories related to the formation of differentiated tissues^9^. Overall, the dynamism of the epigenome suggests a highly concerted organization of DNA structure which first drives cell movement and then the development of the animal body plan.

### Genome-wide analyses implicate key transcription factor motifs involved in developmental lineage decision-making

Having identified genome-wide patterns of chromatin accessibility dynamics and the biological processes associated with them, we next wanted to investigate molecular regulators that might underlie these processes. Thus, we utilized our data to identify patterns of transcription factor (TF) motif accessibility and binding across *Hofstenia* development, reasoning that the most drastic changes likely correspond to TFs that drive gene expression and play a major role in the biological processes at each of these timepoints.

We used chromVAR to investigate the accessibility of motifs genome-wide^20^. We considered a curated set of *Hofstenia* motifs identified in previous studies, which were compiled based on *de novo* motif identification and collapsed redundant motifs that we are currently unable to distinguish between in *Hofstenia*^13^. Similar to our observation that replicates from a given time point clustered together in DiffBind based on peak accessibility, we found that they also clustered together when considering motif accessibility (Figure 3A). The top five motifs with the most variable accessibility across time in our dataset were MSX1, RUNT, HOXD1, ZIC4, and GLI2 (Figure 3B). Notably, although the EGR motif had exhibited the greatest dynamism in a similar analysis during regeneration^11^, we found it exhibited low variability genome-wide in development. Further, we found that one of most dynamic motifs, RUNT, exhibited highest motif accessibility at the gastrula stage and an overall pattern of decreasing accessibility as development progressed (Figure 3C). Of note, during regeneration in *Hofstenia* RUNT exhibited a relatively low chromatin opening index score^11^. In addition to these motifs with high accessibility early in development, we observed motifs with other patterns, such as NFIC, which exhibited increasing accessibility at the post-dimple (Figure 3D), and TCF4, which had low accessibility early but increased over development (Figure 3E). In considering the accessibility of all motifs (Supplement 4B), we found that many of them had low variability in accessibility throughout development. However, when considering these motifs individually, we saw that they also exhibited patterns of variable accessibility, but at a smaller scale. For example, EGR showed higher accessibility at gastrula and pigmented prehatchling than at dimple or post-dimple (Figure 3F).

**Figure 3.**
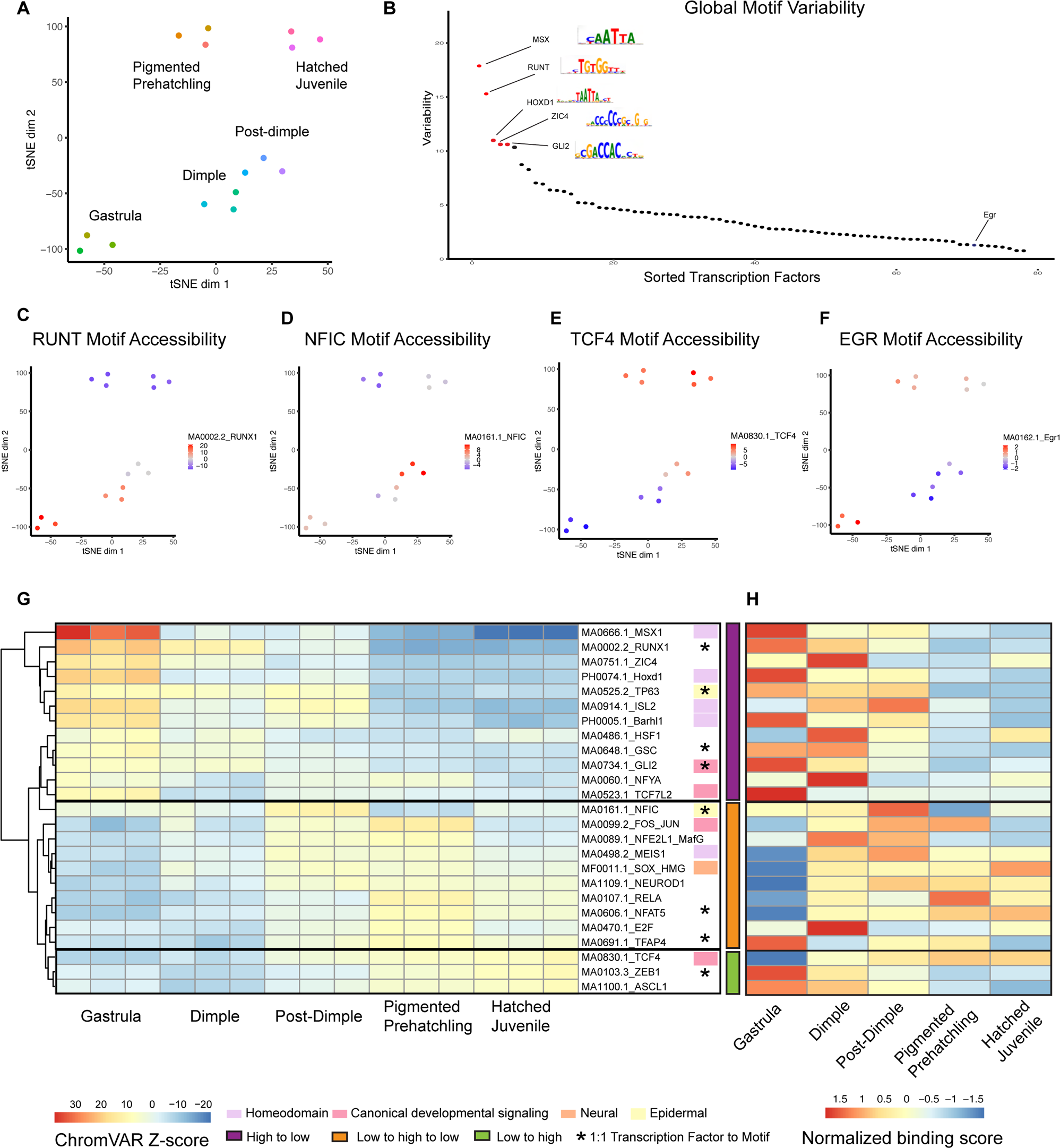
Genome-wide analyses implicate key transcription factor motifs involved in developmental decision making. (A) t-Distributed stochastic neighbor embedding (t-SNE) plot showing groupings of samples, including biological replicates, based on chromatin accessibility of the consensus peak set. Replicate samples are colored on the basis of their time point. (B) chromVAR chromatin variability scores for a curated list of 78 *Hofstenia* transcription factor binding motifs across all time points of development. The top five most variable motifs (Mxs/Hox, Runt, Hoxd1, Zic4 and Gli2) in red, and the most variable regeneration motif (EGR) labeled in blue. (C) Accessibility of RUNT binding motif overlaid on the same t-SNE plot as (A), with scale indicating more accessible chromatin (red) at sites genome-wise in gastrula, dimple, and post-dimple and less accessible chromatin in pigmented prehatchling and hatched juvenile (blue). Note the scale varies for each ChromVar plot. (D) Accessibility of NFIC binding motifs. (E) Accessibility of TCF4 binding motifs. (F) Accessibility of EGR binding motifs. (G) Heatmap of the top 25 most variable motifs at various developmental stages. Motifs with homeodomain in lilac, canonical developmental signaling in pink, neural domains in orange, epidermal in yellow. Three major patterns were colored, high to low (purple), low to high to low (orange), and low to high (green). (H) TOBIAS normalized binding score for each corresponding motif.

When we considered the top 25 most variable motifs, we noted that three main patterns emerged: motifs high at gastrula with decreasing accessibility, motifs that exhibited highest accessibility at post-dimple and/or pigmented prehatchling, and motifs with accessibility that increased progressively until the hatched juvenile time point (Figure 3G). Many of the motifs with highest accessibility corresponded to homeodomain-containing transcription factors (e.g. MSX1, Hoxd1, ISL2, and Barhl1) and showed greatest accessibility at the gastrula stage, potentially pointing to an early role in establishing regions in the animal. We also noted multiple motifs that corresponded to transcription factors implicated in epidermal differentiation during regeneration in *Hofstenia* (e.g. TP63 and NFIC)^21^. The high accessibility of these motifs early in development is consistent with the fact that epidermis is one of the first molecularly discernable lineages^14^. In the group of motifs that exhibit highest accessibility at the post-dimple or pigmented prehatchling stage, we noted a number of motifs that correspond to genes that have known roles in neurogenesis in either *Hofstenia* or other organisms, which is concordant with past work where differentiated neurons appear at pigmented prehatchling^14^. We also found that motifs corresponding to canonical developmental signaling pathways showed variable patterns of motif accessibility. For example, TCF4 and TCF7L2 are both downstream effectors of canonical Wnt signaling, but TCF4 shows greatest accessibility at hatching, while TCF7l2 has the greatest accessibility at the gastrula stage. This could correspond to differential roles for Wnt signaling and utilization of different Wnt ligands at various times during development. However, because there are multiple *Hofstenia* transcripts that correspond to TCF genes, it is difficult to disentangle the relationship individual TCFs might have with patterns of motif accessibility or binding with these data.

We therefore reasoned that focusing on motifs that can be assigned 1:1 to a single *Hofstenia* transcription factor gene might facilitate the use of these data to provide insight into biological processes. To identify such motifs with putative correspondence to a single *Hofstenia* transcript, we found the coding sequence that corresponded to the transcriptional regulator of the JASPAR motif sequence and performed BLAST against the *Hofstenia* transcriptome. We then took each of these transcripts and performed BLAST against the whole NCBI database to confirm this putative identity, and also used the NCBI Conserved Domain Search to confirm that the DNA binding domains associated with that particular TF were predicted to be present in the *Hofstenia* transcript based on nucleotide sequence. We considered genes to be 1:1 with a motif if they occurred as a single copy and could be assigned to one motif.

To more directly predict the presence and binding of motifs by a TF in regions of accessible chromatin, we used the footprinting program TOBIAS. Footprinting analyses take advantage of the fact that TFs bound to DNA hinder cleavage in those small regions, leading to short stretches of decreased signal in regions of otherwise high accessibility. Such regions associated with motifs are predicted to be bound^22^. We uncovered many of the same global patterns, such as highest binding at gastrula for the motifs that have high accessibility at gastrula (Figure 3H, Supplement 4C). When we compared binding and accessibility for a given motif, there was general concordance, such as MSX1 having highest motif accessibility and predicted binding at early time points then decreasing later in development. However, we also noted differences. For example, ZEB1 is predicted to have high binding genome-wide early but high accessibility later in development. This shows that an accessible peak containing a given motif does not necessarily mean a TF is bound, and accessibility and binding can be studied as independent variables.

### RUNT is among the most variably accessible motifs across *Hofstenia* development, suggesting Runt as a major regulator of early development

Of the 1:1 transcription factor (TF) motifs paired with single *Hofstenia* genes, RUNT emerged as one of the most dynamic motifs. To generate hypotheses for when and where the gene *runt* is expressed we revisited developmental single-cell RNA-seq data using the trajectory inference tool URD, which computationally predicts molecular trajectories and allows for the identification of when different groups of cells become transcriptionally distinct from one another^23^. Armed with previous knowledge of which tissue types these branches would give rise to (Figure 4A) we projected the expression of *runt* onto the URD tree^14^. We found that *runt* expression occurs in a number of different branches that lead to a variety of differentiated tissue types where the color of the branch represents the average expression of a cell in pseudotime (Figure 4B). *Runt* expression (Figure 4C) mirrors the pattern of RUNT motif accessibility (Figure 3C), with high levels of *runt* early and decreasing levels as development progresses. Additionally, the number of RUNT sites that TOBIAS identified as putatively bound decreased from gastrula (∼25%) to pigmented prehatchling (∼11%) (Figure 4D).

**Figure 4.**
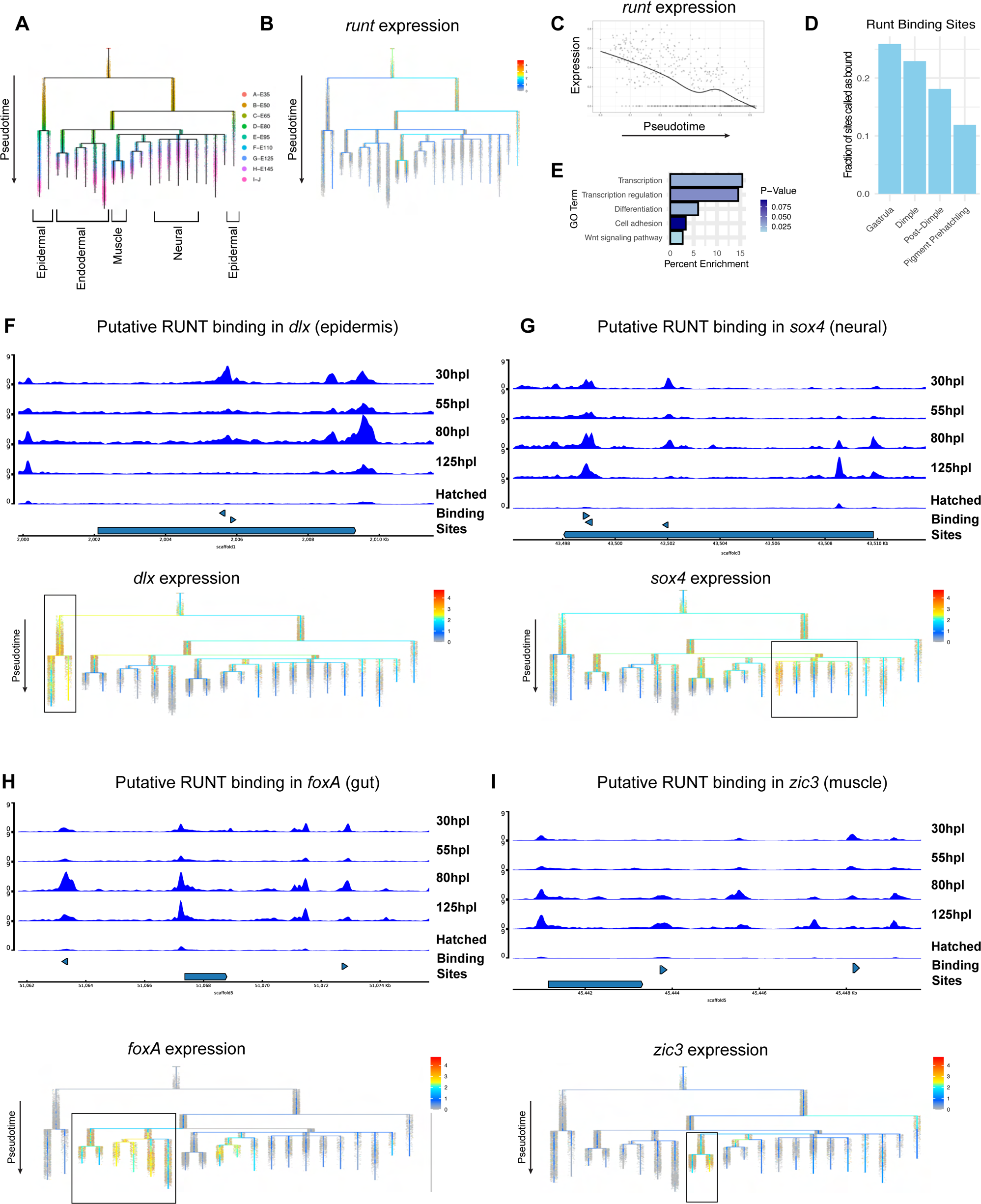
RUNT is among the most variably accessible motifs across *Hofstenia* development, suggesting Runt as a major regulator of early development. (A) URD tree with major tissue lineages labeled (Kimura et al. 2022). (B) URD tree with expression of *runt.* (C) *Runt* expression over URD psuedotime. (D) Fraction of bound RUNT sites. (E) GO Term enrichment for genes with RUNT binding. (F) Accessibility of the *dlx* gene locus with candidate RUNT binding sites. URD tree with expression of *dlx,* epidermal lineage is boxed. (G) Accessibility of the *sox4* gene locus with candidate RUNT binding sites. URD tree with expression of *sox4*, neural lineage is boxed. (I) Accessibility of the *foxA* gene locus with candidate RUNT binding sites. URD tree with expression of *foxA*, gut/endodermal lineage is boxed. (H) Accessibility of the *zic3* gene locus with candidate RUNT binding sites. URD tree with expression of *zic3*, muscle lineage is boxed.

To understand which types of genes might be under Runt control at the earliest timepoints in development we first took peaks regions containing RUNT motifs that TOBIAS had predicted as bound. After linking these peak regions to genes with UROPA, we used GO to characterize which processes these genes may be participating in and found an enrichment for GO terms associated with transcription. To gain higher resolution into what was driving this GO term, we looked at the *Hofstenia* transcripts that contributed to this enrichment (Figure 4E). These genes putatively bound by Runt included several transcription factors that had been previously validated as functional regulators of cell type differentiation during *Hofstenia* regeneration^13,21^. These included the epidermal TF *dlx* (Figure 4F), neural TF *sox4* (Figure 4G), and endodermal/gut TF *foxA* (Figure 4H), which all had binding sites for RUNT. The TF *zic3* appeared in our list of genes with binding sites for RUNT (Figure 4I) and while the expression of this gene had not previously been studied, we found *zic3* expression in the muscle branch of the URD tree (Figure 4I). Notably, all of these genomic loci contained regions of accessible chromatin with predicted binding at a RUNT motif during the gastrula stage. Together these data suggest that Runt may play an early role in *Hofstenia* development and could be responsible for initiating the specification of many major tissue lineages in *Hofstenia*.

Our findings in development are reminiscent of those in the process of regeneration – in previous work investigating *Hofstenia* regeneration, *runt* was characterized as a wound-induced gene, and in examining previous data we found that the loci of *dlx, sox4, foxA*, and *zic3* all carrying predicted RUNT sites bound at six hours post amputation^11^. This suggests that amputations in adult animals could be resulting in rapid upregulation of an early developmental program.

### Motif accessibility and binding provide hypotheses for regulatory relationships of known epidermal differentiation genes

Given that RUNT binding occurred early, but appeared to control many lineage-specific genes, we next focused on transcription factors that have enrichment in specific tissue lineages. We asked whether our ATAC-seq data would corroborate what was already known during *Hofstenia* regeneration about TFs which specify epidermis, *dlx, p73, nfic* and as well as the genes which mark differentiated epidermis, *cah6* and *epiA* ^21,24^. In addition to identifying a putative RUNT motif in the locus of *dlx*, a regulator of epidermal differentiation, we also noted predicted binding of Runt in the genomic locus of another known marker of epidermal progenitors during regeneration, *p73* (Figure 5A)^21^. Given the fact that the functional relationship between epidermal transcription factors and differentiated markers had previously been clarified in Hofstenia regeneration we wanted to ask if there was evidence that the same regulatory linkages between epidermal transcription factors and differentiated factors are maintained in development. We found *p73* mRNA was enriched in the branches leading to epidermis in the URD tree (Figure 5B) and was highly expressed early in development (Figure 5C, 5G). This was consistent with both the chromatin accessibility and predicted binding of TP63 motifs genome-wide (Figure 3G), which likely have a 1:1 association with *p73* in *Hofstenia* (p73 is the only homolog of the p53/63/73 family in *Hofstenia*). An additional known epidermal progenitor marker, *nfic*, also contains TP63 binding sites (Figure 5D)^21^. *nfic* was also expressed and enriched in epidermal lineages during development (Figure 5E, F). This included both the epidermal tissue that covers the periphery of the animal as well as the epidermal lining of the pharynx (Figure 5J). Next, we asked if there was evidence of NFIC binding sites in peak regions associated with the previously characterized *Hofstenia* epidermal genes *epiA* (Figure 5M) and *cah6*^21,24^. Both loci contained putatively bound NFIC motifs (Figure 5I and L), and further, both genes were coexpressed with *nfic* in the predicted epidermal lineages (Figure 5H, K).

**Figure 5.**
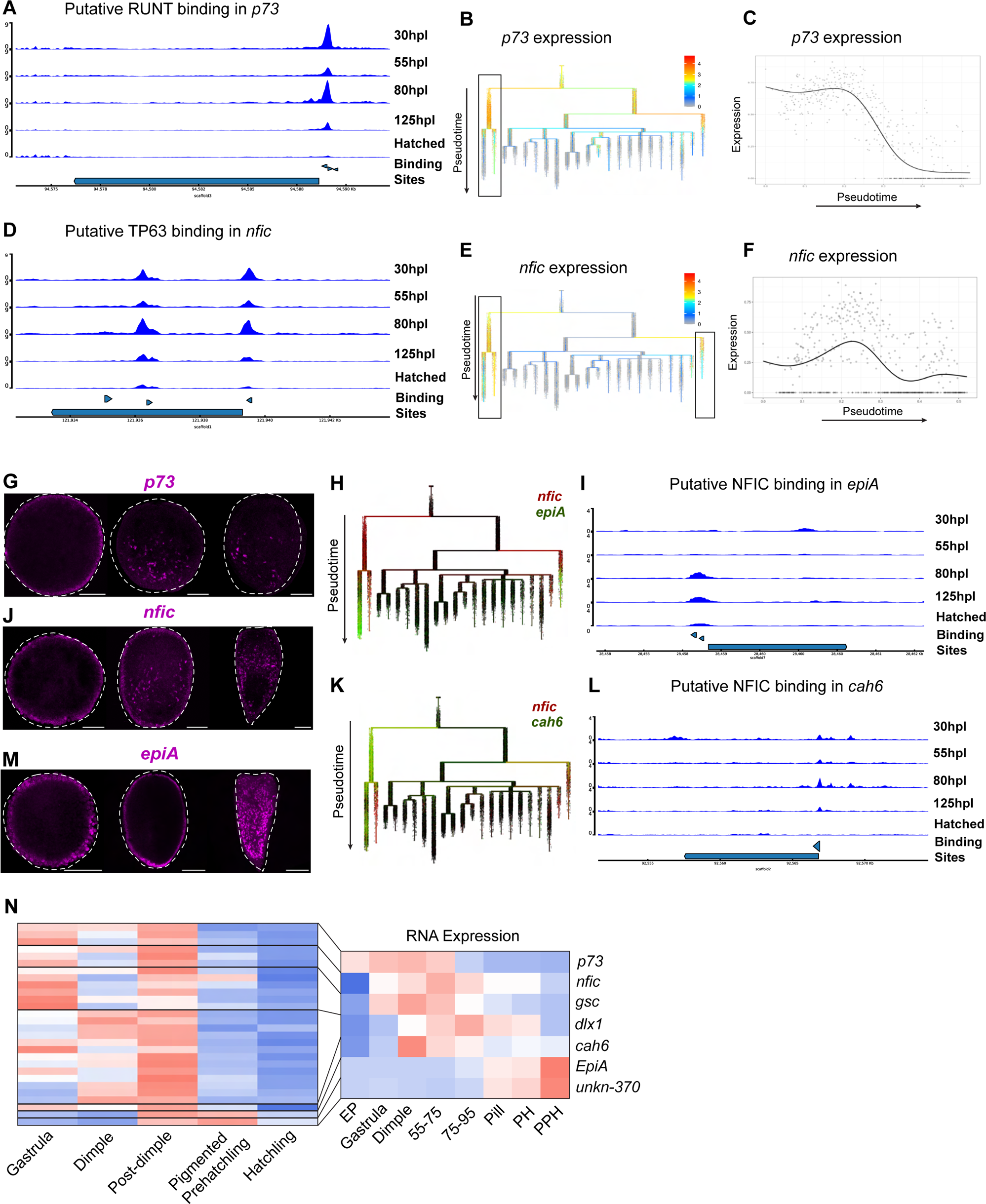
Motif accessibility and binding provide hypotheses for regulatory relationships of known epidermal differentiation genes. (A) Accessibility of the *p73* gene locus with candidate RUNT binding sites. (B) URD tree with expression of *p73,* epidermal lineage is boxed. (C) *p73* expression. (D) Accessibility of the *nfic* gene locus with candidate TP63 binding sites. (E) URD tree with expression of *p73,* epidermal lineage, along with pharyngeal epidermis is boxed. (F) *nfic* expression (G) *in-situ* gene expression of *p73* at dimple, post-dimple, and pigmented prehatchling (H) URD tree with the coexpression of *nfic* in red and *epiA* in green. (I) Accessibility of the *epiA* gene locus with candidate NFIC binding sites. (J) *in-situ* gene expression of *nfic* at 55hpl, 80hpl, and 125hpl. (K) URD tree with the coexpression of *nfic* in red and *cah6* in green. (L) Accessibility of *cah6* gene locus with candidate NFIC binding sites. (M) *in-situ* gene expression of *epiA* at dimple, post-dimple, and pigmented prehatchling. (N) Linked ATACseq peak-accessibility and corresponding RNA expression of previously described epidermal markers. Abbreviations are EP (Early Pooled), PH (Prehatchling), PPH (Pigmented Prehatchling).

Overall, the developmental ATAC-seq data corroborate that the epidermal differentiation trajectory first uncovered in homeostasis and regeneration holds true during embryonic development. Further, these data provide hypotheses for regulatory relationships between these genes and implicate Runt as an upstream regulator of this differentiation trajectory during development. These data also allowed us to examine putative cis-regulatory elements by linking broad patterns of peak accessibility to genes and corresponding RNA levels, i.e. gene expression (Figure 5N). Corresponding to the early emergence of the epidermal lineage in the single-cell RNA-seq data, peaks associated with epidermal genes became accessible early in development. Further, peaks associated with a gene tended to become accessible prior to that gene’s expression, although there is some variability in peak accessibility dynamics even in peaks associated with the same genes - potentially representing finer regulation of gene expression such as with enhancer or repressor elements.

### Predicted transcription factor binding uncovers a regulatory network with Pitx, Fox, and Zic transcription factors in *Hofstenia* muscle development

Given the impact of developmental chromatin profiling data in corroborating known differentiation trajectories, we next sought to test the power of these data to reveal new mechanisms of differentiation during *Hofstenia* development. Our analysis of RUNT motifs had revealed putative binding in *zic3*, a previously undescribed *Hofstenia* transcription factor (TF) that we found is enriched within the muscle lineages (Figure 4H). Prior work in *Hofstenia* muscle development focused on differentiated muscle markers, not on developmental regulators. Broadly, muscle cells appear around 80hpl, late in the post-dimple stage, followed by elaboration of muscle structures until all juvenile muscle types are formed at the time of hatching (pharynx, parenchymal muscle, body wall muscle, and peripheral muscle)^9,24^ (Fig 6B).

**Figure 6.**
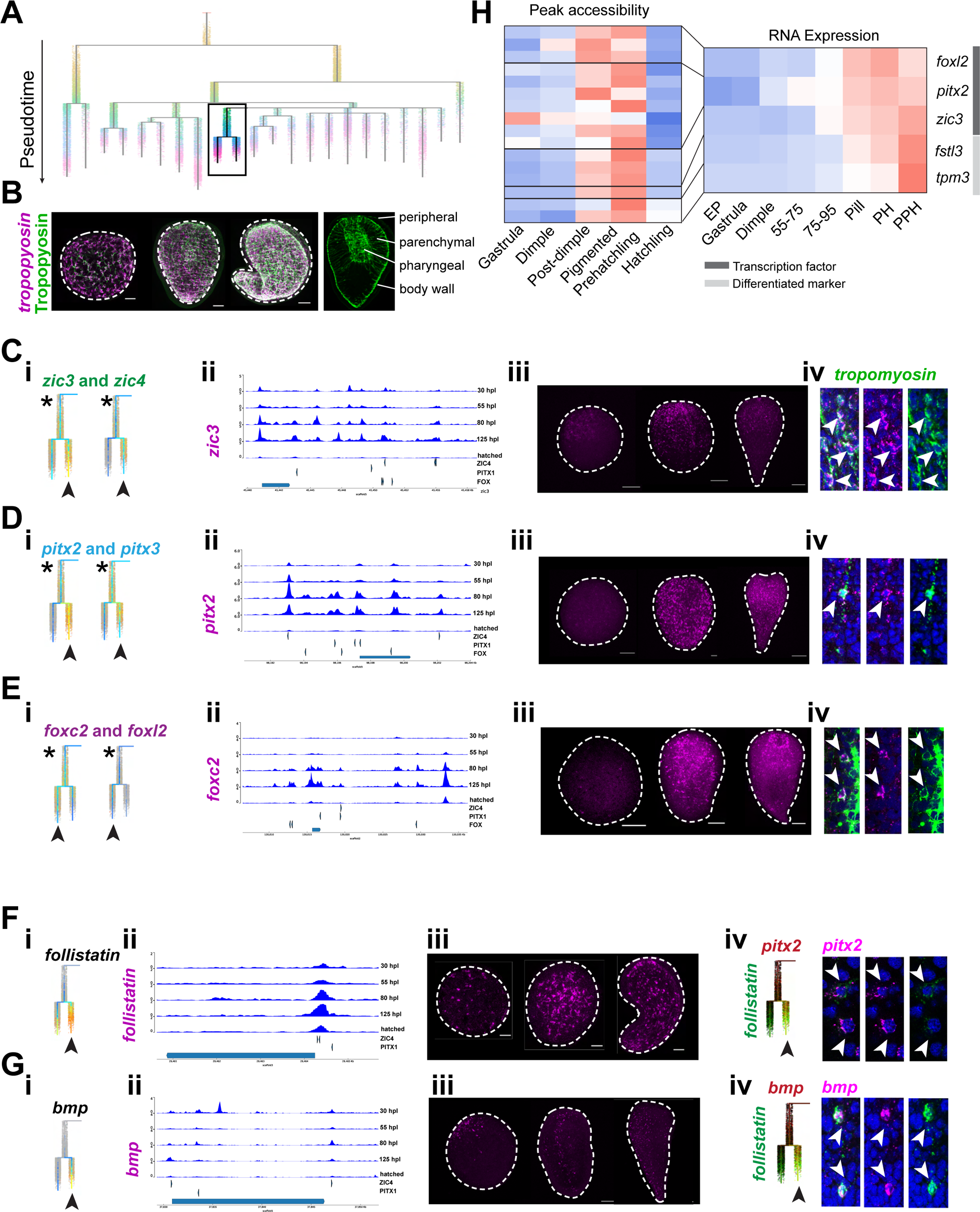
Predicted transcription factor binding uncovers a regulatory network with Pitx, Fox, and Zic transcription factors in *Hofstenia* muscle development. (A) URD tree with muscle lineages boxed. (B) *tropomyosin* gene expression (purple), and tropomyosin antibody stain in developing *Hofstenia* embryos. Peripheral, parenchymal, pharyngeal, and body wall muscles are labeled. (C-E) (i) URD tree expression of *zic3* and *zic4; pitx2* and *pitx3*; or *foxc2* and *foxl2* in the muscle branches, (ii) accessibility of *zic3, pitx2*, and *foxc2* loci with candidate ZIC4, PITX1, and FOX binding sites, (iii) *in-situ* gene expression of *zic3, pitx2*, and *foxc2,* and (iv) coexpression of *zic3, pitx2*, and *foxc2* with *tropomyosin*, white arrows denote coexpression. (F) (i) URD tree expression of *follistatin*, (ii) accessibility of *follistatin* locus with candidate ZIC4, PITX1 binding sites, (iii) *in-situ* gene expression of *follistatin*, and (iv) coexpression of *follistatin* with *pitx2*. (G) (i) URD tree expression of *bmp*, (ii) accessibility of *bmp* locus with candidate ZIC4, PITX1 binding sites, (iii) *in-situ* gene expression of *bmp*, and (iv) coexpression of *bmp* with *follistatin*. (I) ATACseq peak-accessibility and corresponding RNA expression of transcription factors and differentiated markers of muscle. Abbreviations are EP (Early Pooled), PH (Prehatchling), PPH (Pigmented Prehatchling).

To investigate whether *zic3* has a role in muscle formation during development, we asked where *zic3* might be binding by screening loci with predicted binding at ZIC4 motifs. Previously, we had investigated motifs that had a predicted 1:1 correspondence with a gene. In contrast, here we found five *zic* genes in the *Hofstenia* transcriptome. Of these, two were enriched in muscle *(zic3 and zic4)* (Figure 6C). One *zic* did not have detectable expression in the single-cell RNA-seq data (*zic2*), and two of the *zics* showed early expression (*zic4-2* and *zic3-2*) (Supplement Figure 6A). Although we cannot firmly distinguish between which of these Zic proteins may bind to different predicted ZIC4 binding sites in the *Hofstenia* genome, we reasoned that by considering predicted binding at later developmental timepoints, after muscle has a distinct transcriptional signature and *zic4-2* and *zic3-2* expression have diminished, in conjunction with genes that are enriched in the muscle URD branches, might provide hypotheses for which genes *zic3* and *zic4* are regulating.

When we asked which genes were associated with putatively bound ZIC4 sites, we identified additional muscle-enriched TFs, including two Fox homologs (*foxc2, foxl2*) and two Pitx homologs (*pitx2 and pitx3*). We noticed that these transcription factors are expressed in the earliest time points in the muscle lineage, in a putative muscle progenitor population (Figures 6C.i, D.i, E.i, asterisks). We thus sought to ask whether these genes might have a role in regulating each other’s transcription. When we considered motifs corresponding to *pitx* (PITX1), *fox* (FOX), and *zic* (ZIC4), we focused on sites with predicted binding in peaks associated with the loci of each of the transcription factors we had identified. This is consistent with regulation between these transcription factors having a role in establishing muscle differentiation (Figures 6C.ii, D.ii, E.ii). We also reasoned that if these genes had a role in muscle differentiation, we might see co-expression with differentiated muscle markers. When we looked at the *Hofstenia* pan-muscle marker *tropomyosin*, we found that although not restricted to *tropomyosin*+ cells, all transcription factors were coexpressed with some *tropomyosin*+ cells (Figures 6C.iv, D.iv, E.iv). We also noted that these transcription factors had different gene expression patterns from each other in the embryos, and none of them included the full domain of *tropomyosin* expression (Figure 6B). While *pitx2* and *pitx3* were peppered throughout the animal, *zic3* and *zic4* were concentrated in the anterior, as were *foxc2* and *foxl2* (Figures 6C.iii, D.iii, E.iii). The variable expression patterns of these TFs made us wonder whether they might not be broad muscle regulators, but rather involved in differentiation or specification of sub-populations of muscle. Following the bifurcation of the muscle lineage at later developmental stages, we saw that the Zic and Pitx genes were enriched in one branch, while *foxc2* and *foxl2* had higher enrichment in the other branch (Figures 6C.i, D.i, E.i, arrows).

Thus, to better understand the downstream roles of these transcription factors in muscle specification, we considered other genes that were bound by motifs predicted to be associated with these transcripts (ZIC4, PITX1, and FOX). We saw genes with predicted structural roles in muscle, such as *titin-15*, additional transcription factors, and a number of genes previously determined to have patterning roles during regeneration and homeostasis, consistent with *Hofstenia* muscle’s dual roles as a contractile tissue and in dispensing patterning information^12,25^. Many of these had expression in subregions of the musculature, such as the mouth or pharynx, supporting the idea that these transcription factors may be involved in specifying subpopulations of muscle (Supplement 6B). One striking example was *follistatin*, which had previously been identified as an early wound-induced gene during regeneration^11^. *follistatin* was also identified as a marker of muscle clusters in *Hofstenia* post-embryonic scRNA-seq data^21^. During development, *follistatin* is enriched in the same muscle branch as the *pitxs* and *zics*, and contains binding sites for both ZIC4 and PITX1 (Figure 6F). Furthermore, we found that some, although not all, *follistatin*+ cells were coexpressed with *pitx2*, consistent with regulation by *pitx2*. When we considered other markers of this branch, we saw that they contained a number of previously identified patterning genes, such as *pk-1, dgo-1, wnt-5,* and *bmp*^10,26^. In support of this observation, we found that *bmp* is co-expressed with *follistatin* (Figure 6G.iv), and further has both ZIC4 and PITX1 binding sites (Figure 6G.ii). We saw that the loci of many additional genes enriched in this branch also contain PITX1 and/or ZIC4 binding sites. This led us to hypothesize that Zic and Pitx transcription factors regulate a *follistatin*+ population of muscle that has patterning roles in the animal. This is reminiscent of a finding in planarian regeneration that different populations of muscle have different instructive roles in regeneration^27^.

Finally, we sought to investigate how these findings fit into known features of muscle development by comparing chromatin accessibility with mRNA expression of muscle markers. When we compared accessibility of peaks associated with these genes and their mRNA expression, we noted that chromatin accessibility associated with a given locus preceded that of the corresponding mRNA expression (Figure 6H). Furthermore, when we considered TFs relative to differentiated markers, we found that expression preceded that of differentiated markers, consistent with the possibility that these TFs might have a role in regulating genes expressed in differentiated muscle cells. The predominant pattern of peak accessibility was that the peaks in the loci of muscle-enriched TFs were high at the post-dimple stage, while differentiated factors showed only a slight increase of peak accessibility at the post-dimple stage, with increased accessibility at pigmented prehatchling. This timing is consistent with the previous descriptions of the timing of *Hofstenia* muscle cell development, where the onset of differentiated muscle marker expression was observed in some cells at the post-dimple, followed by a more detectable fibrous muscle cell morphology at later time points, correspondent with an increase in mRNA expression (Figure 6B)^9,24^. Notably, we observe a similar temporal relationship between ATAC-seq peaks and mRNA, as well as between TFs and differentiated factors, in both epidermal and muscle lineages, even though these tissues are specified at very different times during development (Figure 6C)

Overall, this demonstrates the power of these data to corroborate our previous knowledge about the timing of muscle development as well as to generate new hypotheses about how subpopulations of muscle are specified, and how patterning information required for the adult animal is established during development. These data allowed us to connect gene expression with chromatin opening for known markers of muscle in development, as well as to develop hypotheses for the regulation of subpopulations of muscle, one of which harbors a number of known patterning genes required for regeneration.

## DISCUSSION

Here, we sought to identify patterns of opening and closing chromatin during embryogenesis in the acoel *Hofstenia miamia* with the objective of creating a large dataset that is powerful for generating hypotheses for developmental mechanisms.

Focusing on the broad patterns in these data across the genome, we found: **1)** the dimple stage was depauperate in terms of accessible chromatin. This lack of active regulatory regions corresponded to a transition in numbers of cells that were positive for active RNA transcription, with very few transcribing cells before the dimple stage and the subsequent stages having transcription in the majority of cells. Therefore, it is possible that the lack of ATAC-seq peaks at the dimple stage indicates that the full maternal-to-zygotic transition occurs late in development, after the dimple stage. **2)** Our studies of epidermal and muscle differentiation showed that increased chromatin accessibility of a gene locus generally precedes transcription of the corresponding mRNA, consistent with published work showing that chromatin opening is necessary albeit not sufficient for gene expression^28^ (**3)** In the stages of development we sampled, ∼90% of identified regulatory regions showed dynamism over time, similar to high proportions of dynamic peaks in crustaceans^2^. This suggests that few processes are stable across developmental time.

In addition to mechanisms operating at a genome-wide scale, we utilized these developmental ATAC-seq data in combination with single-cell RNA-seq data to generate hypotheses for regulatory linkages for differentiation processes in *Hofstenia*: **1)** Runt emerged as an early regulator, putatively needed for the activation of transcription factors for differentiation of multiple tissues, including epidermis, muscle, neurons, and gut. Given that Runt homologs play important roles in early development in other organisms^29^, functional studies are needed to make specific comparisons of how Runt’s role in *Hofstenia* compares to its role in other species. For example, in a related symbiotic acoel species *Convolutriloba longifissura*, *runt* is induced after injury and is required for regeneration but also affects the algal transcriptional response^30^. **2)** We found that regulators of epidermal differentiation known from studies of homeostasis and regeneration likely play the same role in development, but further, our data provided regulatory linkages for these genes that can be functional tested in *Hofstenia* given the availability of tools such as genome-editing and transgenesis. **3)** Our data also yielded novel developmental mechanisms, uncovering a putative gene regulatory network (GRN) for the formation of two distinct muscle populations in *Hofstenia*. Given the importance of muscle as the source of patterning information during regeneration in *Hofstenia* and planarians^25,27^, functional probing of this GRN will reveal how patterning information is restricted to muscle cells. Further, the involvement of Zic, Pitx, FoxC, and FoxL homologs in this GRN indicates conservation of muscle differentiation programs across animals, given the known functions of these genes in muscle development in, echinoderms, urochordates, and vertebrates^31–33^.

Overall, we provide here new insight into acoel development as well as a dataset that can be mined by others to identify new developmental mechanisms. Given the unique features of *Hofstenia* as a research organism, i.e. that it enables studies of the entire processes of whole-body regeneration and of embryogenesis, the data generated here will also be powerful for informing comparisons of regeneration and development. Further, given the phylogenetic position of xenacoelomorphs as deep-branching bilaterians, these mechanisms will also inform the evolution of development.

## METHODS

### Immunohistochemistry

Embryos were dechlorinated according to Kimura 2022 and were fixed in 4% paraformaldehyde in artificial seawater (ASW) overnight at 4°C, then washed three times with PBST. Embryos were blocked with goat serum for 1hr then incubated in primary antibody overnight at 4°C. Primary antibody used was mAbs H5 a mouse IgM monoclonal antibody that binds a distinct phosphorylated epitope on the carboxy-terminal domain (CTD) of the large subunit of RNA polymerase II^34–36^. The next day three washes in PBST were performed before a secondary antibody, BODIPY-phallicidin, and Hoescht were added. After incubation overnight at 4°C, three washes were performed before mounting and imaging. Custom *Hofstenia* Tropomyosin antibodies were used as previously described^37^ following dechorionation.

### Generation of ATAC-seq data

ATAC-seq was performed according to standard protocols^38,39^ with minor modifications. Pooled embryos were isolated and placed directly into 200 μL of ice-cold lysis buffer and manually pipetted until a homogeneous solution was obtained. The solution was then filtered with a 40 μm cell strainer to obtain a single cell solution which was then processed as a standard ATAC experiment. All centrifugation steps were carried out at 500 xg. Amplified libraries were run on an Agilent Tapestation 2200 (Agilent Technologies) using a D5000 DNA ScreenTape to assess quality by visualizing nucleosomal laddering. Biological replicates were performed in triplicate for all ATAC experiments. Paired-end 75-bp reads were obtained using an Illumina NextSeq system.

### Sequence Processing

Demultiplexed reads were trimmed of adapters using NGmerge^40^. Trimmed reads were aligned to the *Hofstenia* genome using Bowtie2^41^ which does not include the mitochondrial genome leaving mitochondrial reads unmapped. PCR duplicates were removed using Picard (http://broadinstitute.github.io/picard/). Peaks were called using Genrich in replicate mode which analyzes each sample individually and calculates *p-* values which combined with Fisher’s method into a new corresponding *q-*value (Gaspar 2018) with a gap size of 10 and a minimum AUC of 50 for a peak.

### Chromatin Analysis with DiffBind, chromVAR, and TOBIAS

#### DiffBind

We used the R program DiffBind to calculate statistically significant differences in peak accessibility between timepoints. To calculate counts, we set min.members = 2 and summits =250 to ensure peaks were called in at least two samples, and to set peaks to a uniform width of 500 base pairs. To identify dynamic peaks during development, we performed all pairwise comparisons between time points. For each comparison, this resulted in a list of peaks that had significantly different accessibility between the two time points. To create a set of all dynamic peaks, we took the resulting peaksets from these comparisons and used BEDtools^42^ to concatenate, sort, and merge these lists of peaks. We then used this peakset as an input into DiffBind, along with all of the samples, to create a table of read normalized counts in these peak regions across all timepoints. These counts were then used in downstream hierarchical clustering and fuzzy-c-means clustering. For fuzzy c-means clustering: A count matrix of ATAC-seq peaks was used as input into the R program Mfuzz^17^. Following standard filtering of null values, the fuzzy-c means soft clustering was used with a fuzzifier value m and cluster number that minimized overlap between clusters.

#### chromVAR

In order to use chromVAR^20^, we first forged a BSgenome package for *Hofstenia miamia* using BSgenome^43^. We utilized chromVAR with default parameters according to a standard walk through (https://greenleaflab.github.io/chromVAR/). We used consensus peaksets derived from DiffBind as inputs, the peaks of which were fixed at the center and standardized to a width of 200 bps. For motif matching, we used a curated *Hofstenia* motif set^13^.

#### TOBIAS

Transcription factor binding analysis and footprinting were performed with TOBIAS following a standard run through. Our curated Hofstenia motif set was used as input^22^.

### Gene Ontology Enrichment

Gene ontology (GO) enrichment analysis was done using the same methods as previously described^9,13^. To determine biological processes associated with the various peak sets we used UROPA^44^ to link regions of the genome to nearby genes. GO was performed using the DAVID tool. Uniprot IDs corresponding to Hofstenia genes^10,11^ were used as input into GO, along with a curated *Hofstenia* background list consisting of all Uniprot IDs found to be associated with a Hofstenia transcriptome.

#### *in situ* hybridization

Embryos were dechorinated according to Kimura et al. 2022 (32 mM sodium hydroxide, 0.5 mg/ml sodium thioglycolate and 1 mg/ml of pronase in artificial seawater) and were fixed in 4% paraformaldehyde in ASW overnight at 4°C and stored either 100% methanol at −20 °C or up to 1 week at 4°C until use. Digoxigenin labeled riboprobes were synthesized as previously described and fluorescence in situ hybridizations (FISH) were performed following the protocol described in Srivastava et al. 2014.

## ACKNOWLEDGEMENTS

We thank the members of the Srivastava Lab for their input, particularly Carlos Rivera-López and Catriona Breen for helpful discussions and input for ATACseq analysis, and Allison Kann for input on immunostaining. We also thank Jonchee Kao for suggesting the H5 antibody in Figure 1. We thank the Bauer Core Facility at Harvard University and the FAS Informatics Group for supporting this work.

