## Supplement 1 for "Chromatin profiling data indicate regulatory mechanisms for differentiation during development in the acoel *Hofstenia miamia*"

#### Fragment Distribution of ATAC-seq Libraries

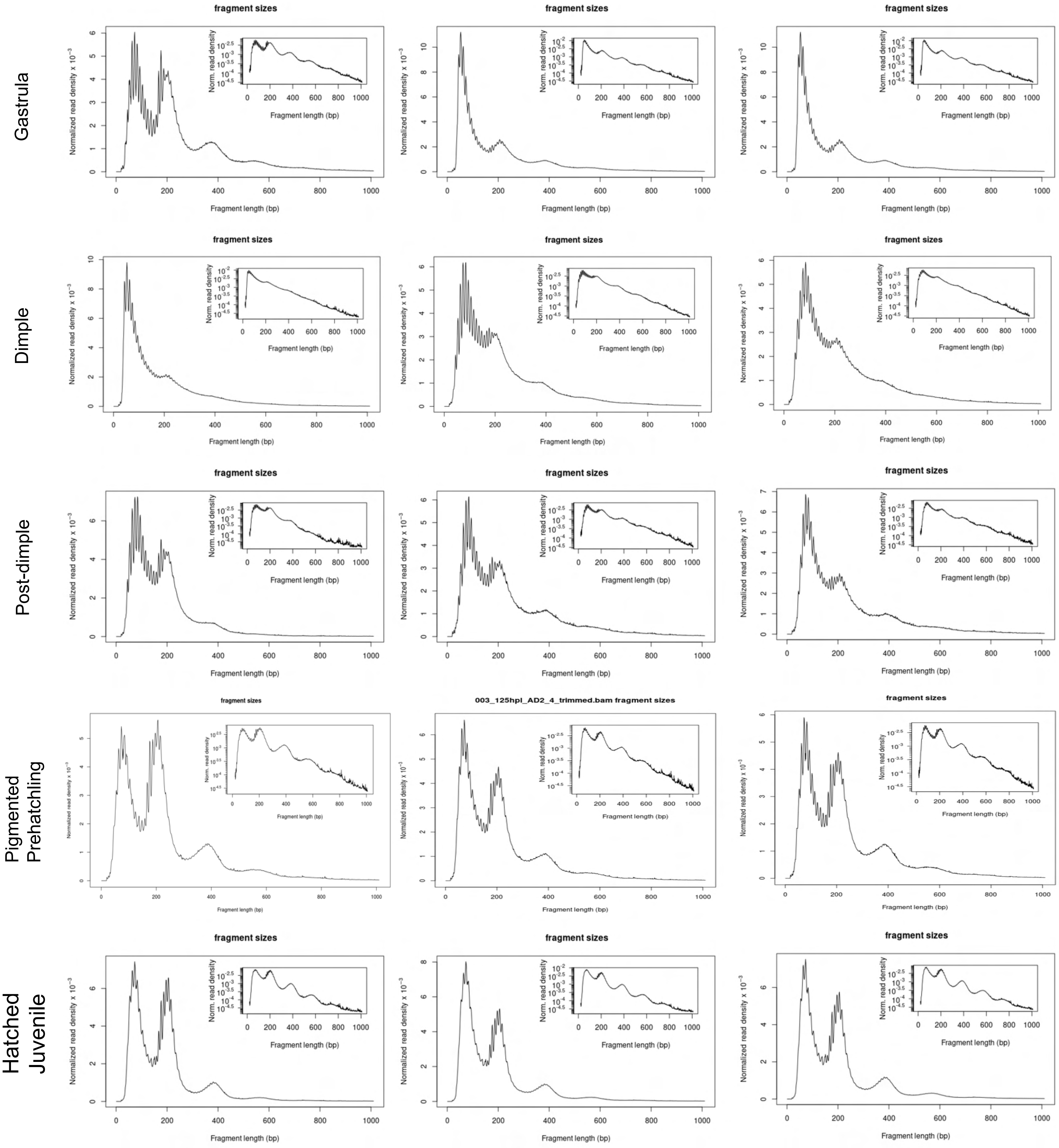

### Supplement 2

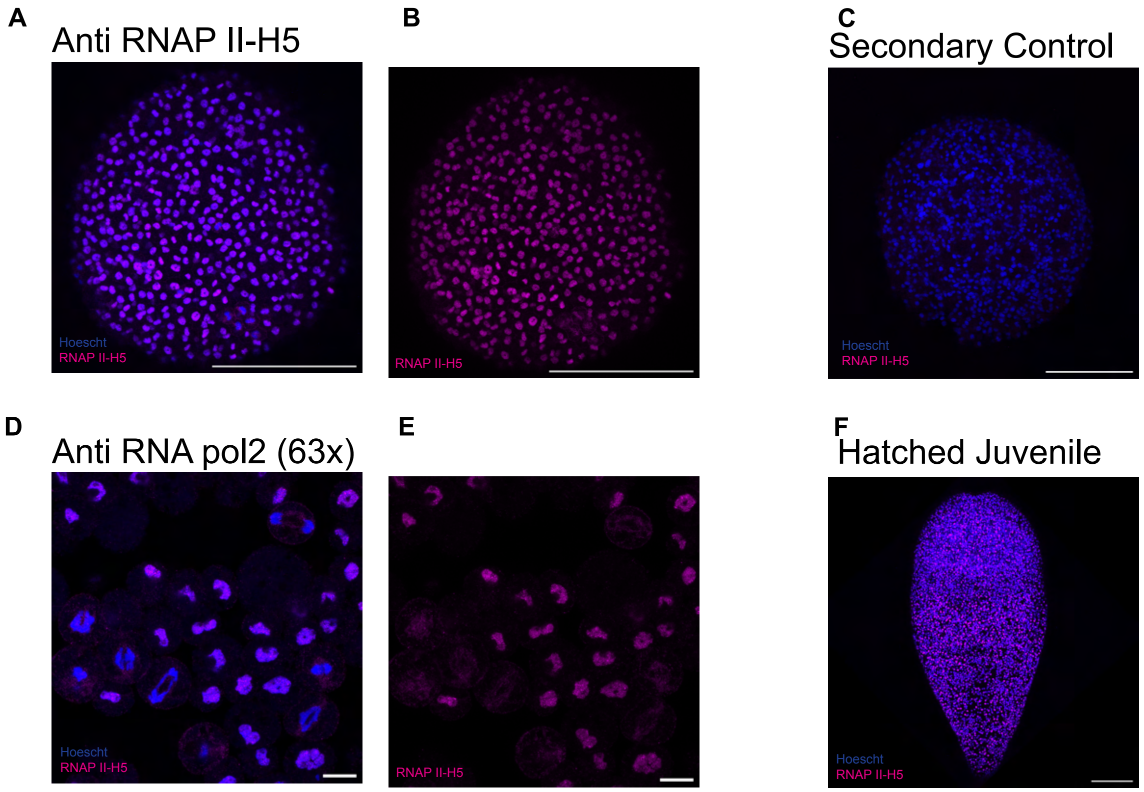

### Supplement 3

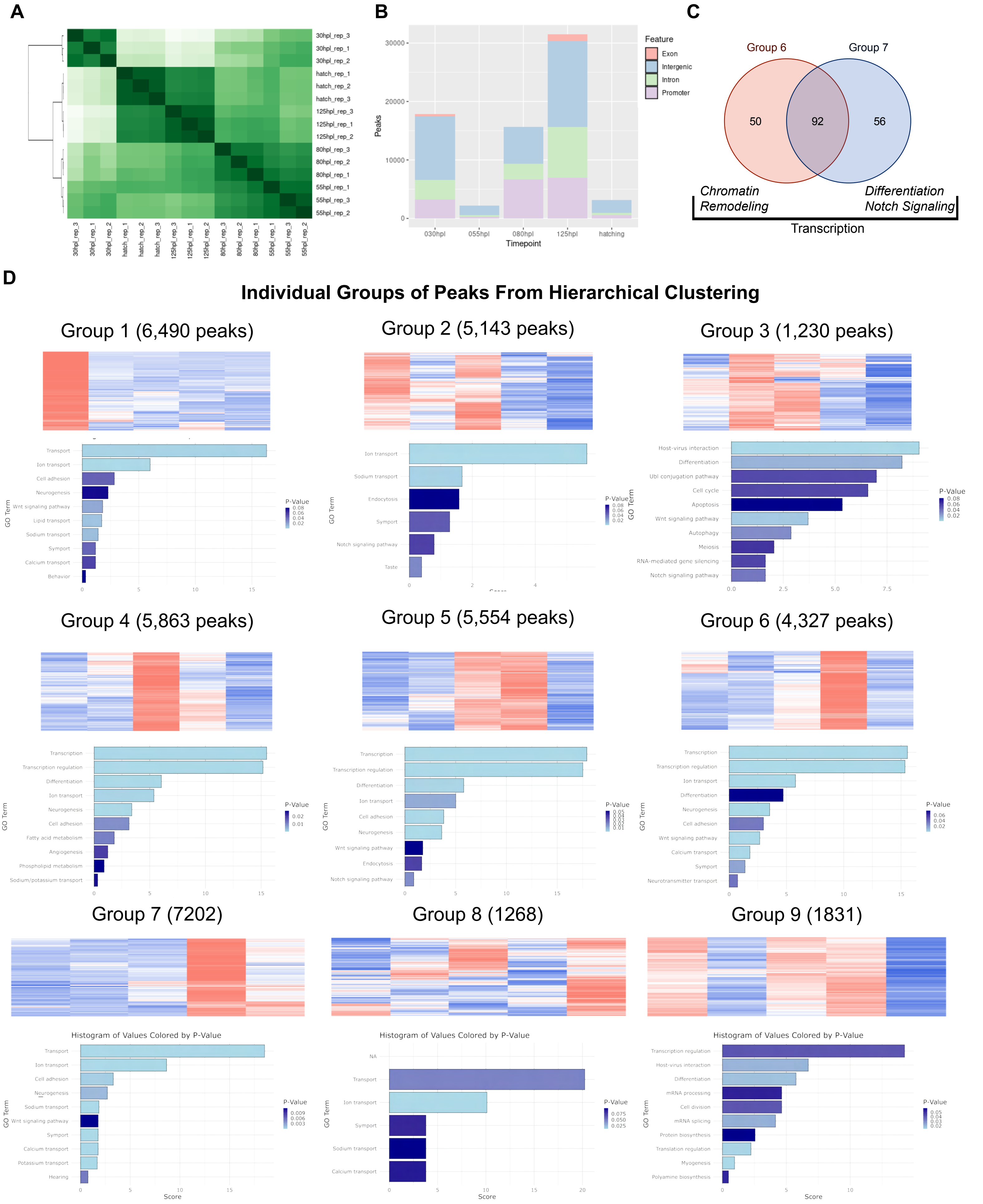

### Supplement 4

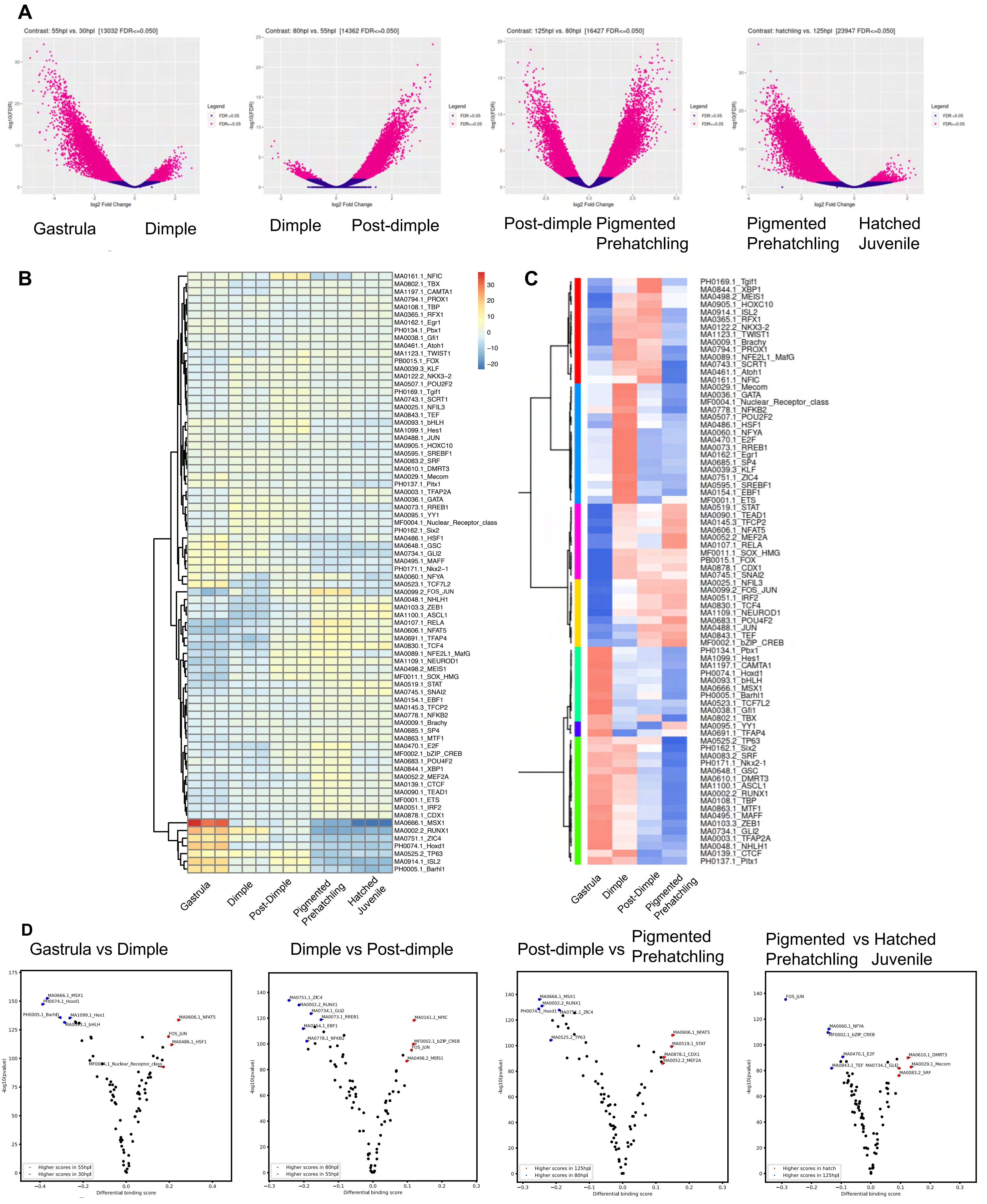

### Supplement 5

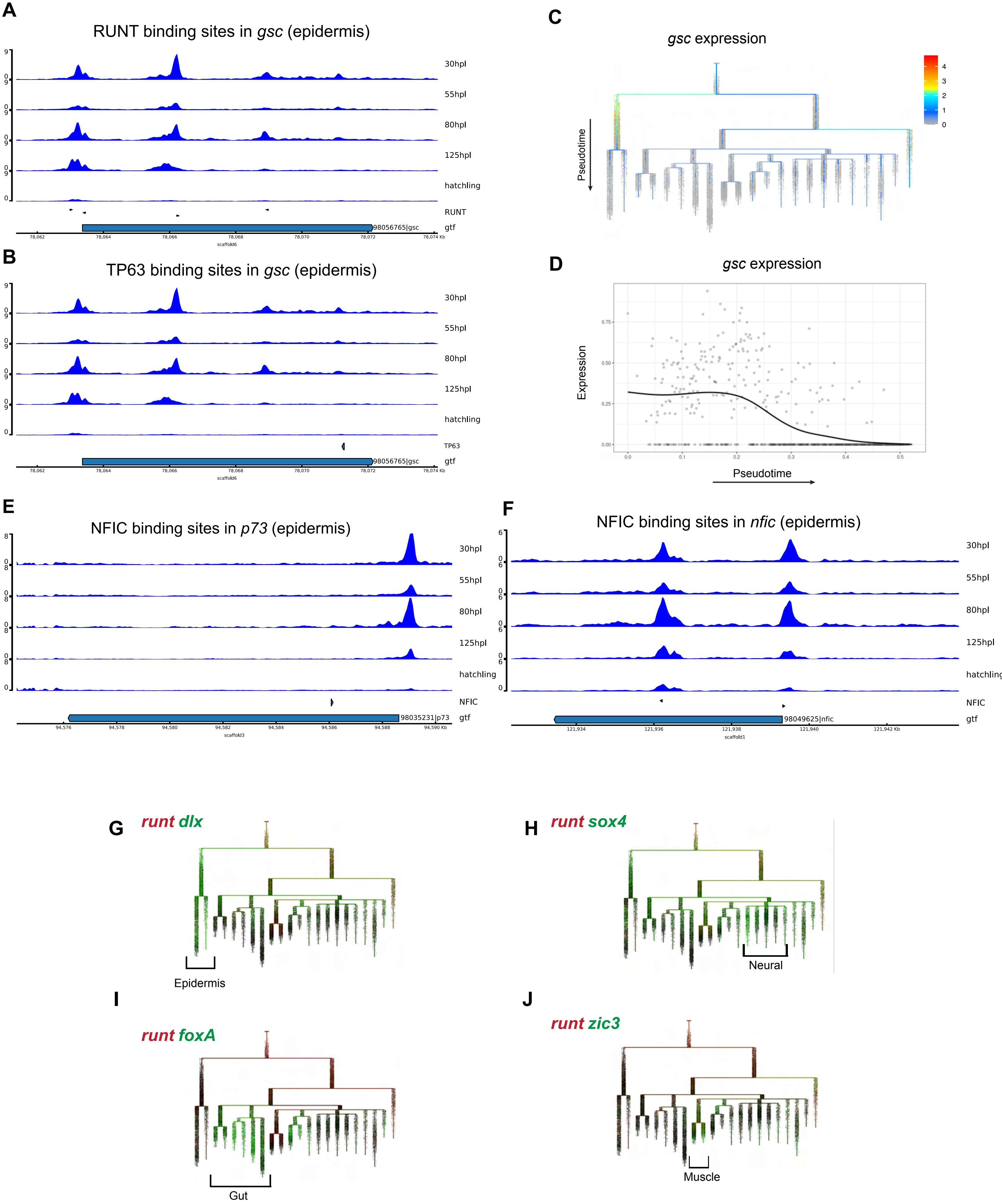

### Supplement 6

**A** Expression of zics in the *Hofstenia* transcriptome on the URD tree

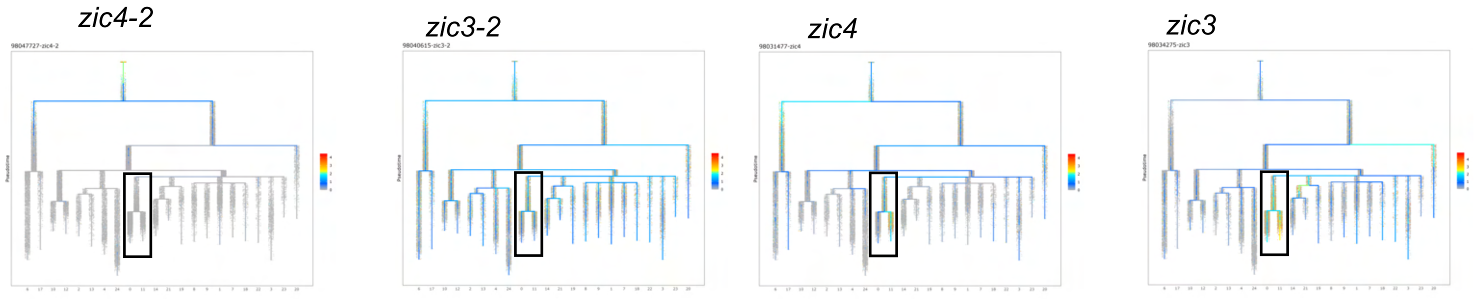


**B**

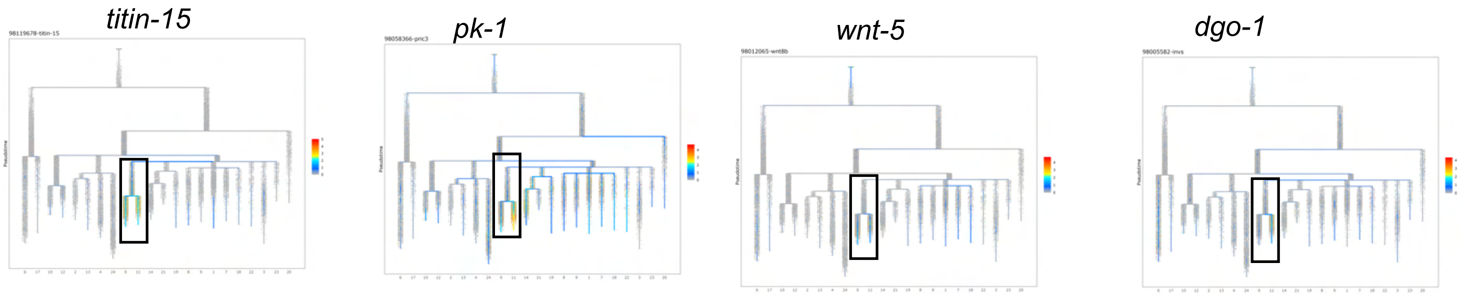


**C**

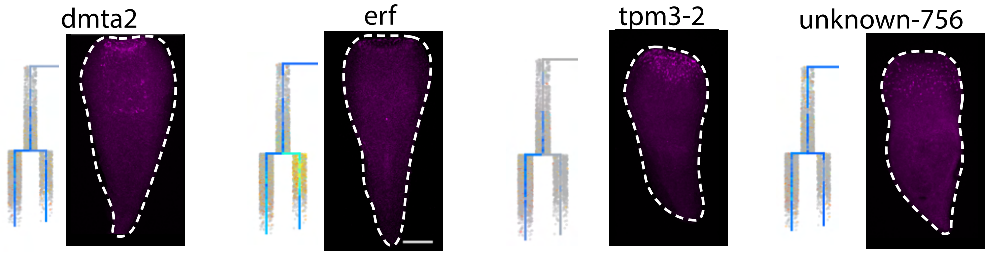
